## Supplementary figures for "β-Catenin condensation facilitates clustering of the cadherin/catenin complex and formation of nascent cell-cell junctions"

### Supplementary figure 1

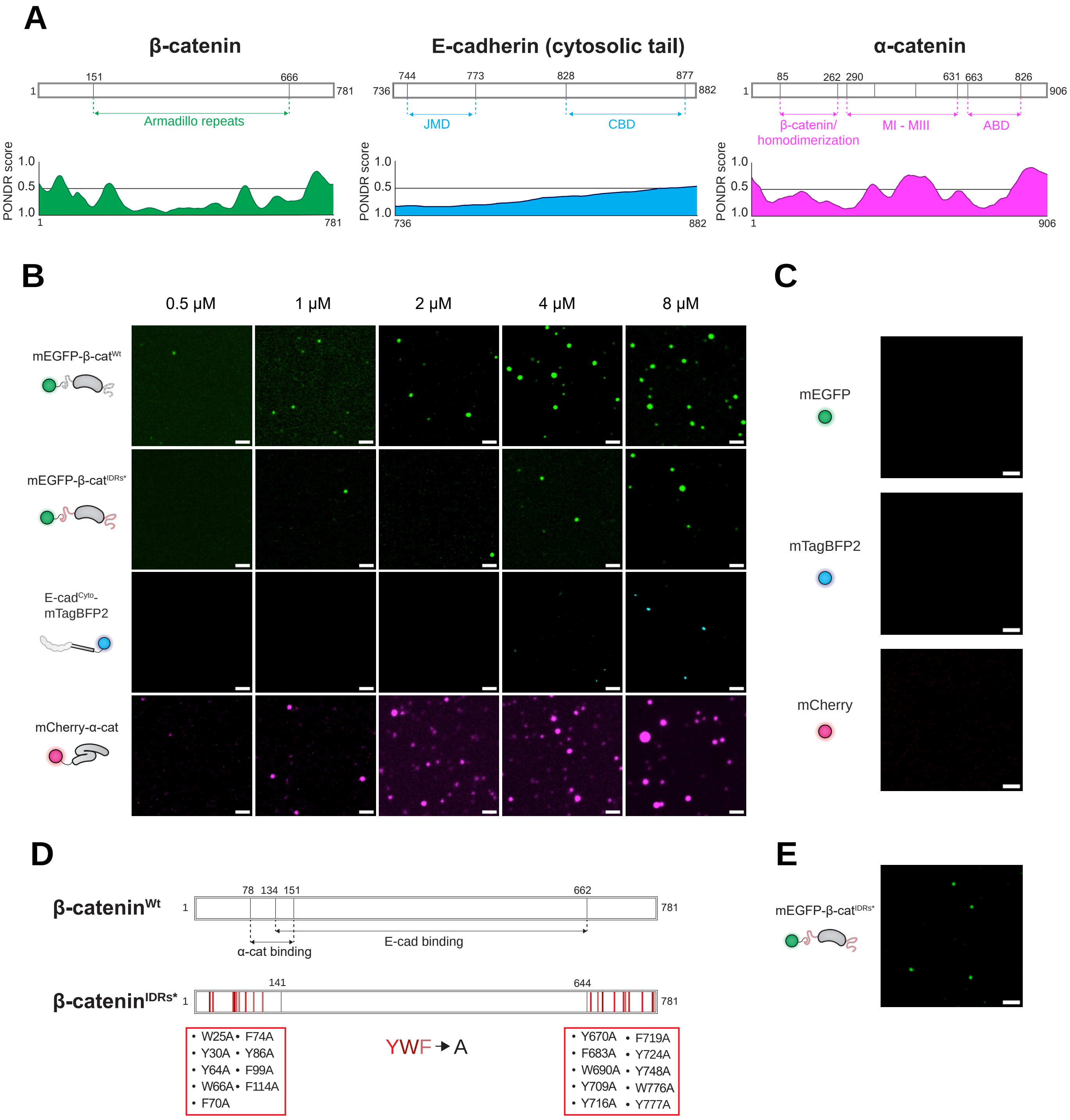


**Figure S1. Ability of E-cadherin$^{Cyto}$, α-catenin, β-catenin$^{Wt}$ and β-catenin$^{IDRs*}$ to form droplets in vitro.**
**A.** Schematic representation of the β-catenin, α-catenin and E-cadherin$^{Cyto}$ domain architectures and their PONDR disorder score predictions. JMD, juxtamembrane domain; CBD, catenin binding domain; MI-MIII, (auto-inhibitory) middle domains; ABD, actin binding domain. **B.** Representative images of droplet assay with purified mEGFP-β-catenin$^{Wt}$, mEGFP-β-catenin$^{IDRs*}$, mCherry-α-catenin and E-cadherin$^{cyto}$-mTagBFP2 at increasing concentrations (0.5 – 8 µM). Note that the contrast of individual images were adapted to properly visualize droplets in all conditions, while the same contrast was used for mEGFP-β-catenin$^{Wt}$ and mEGFP-β-catenin$^{IDRs*}$ at similar concentrations. **C.** Representative image of the purified control fluorophores mEGFP, mCherry and mTagBFP2 at 1 µM. **D.** Schematic representation of β-catenin, indicating the regions involved in its interaction with E-cadherin (E-cad) and α-catenin (α-cat) (top), and the mutated residues of the IDRs* mutant (bottom). E-cadherin binds the Armadillo domain of β-catenin (amino acids 134-662)[39–41], whereas the interaction with α-catenin involves one α-helix in the N-terminal region proximal to the Armadillo domain (amino acids 120-141) and an additional N-terminal helix (amino acids 85-98)[41,58]. In the β-catenin$^{IDRs*}$ mutant 19 single amino acid substitutions are introduced in the N- and C-terminal intrinsically disordered regions (nIDR and cIDR, respectively), replacing tyrosines (T), tryptophans (Y) and phenylalanines (F) by alanines (A). **E.** Representative images of droplet assay of purified mEGFP-β-catenin$^{IDRs*}$ (in the presence of unconjugated mCherry and mTagBFP2, not shown). The proteins were premixed at a concentration of 1 µM before the addition of 10% PEG-8000. All scale bars represent 2 µm.

### Supplementary Figure 2

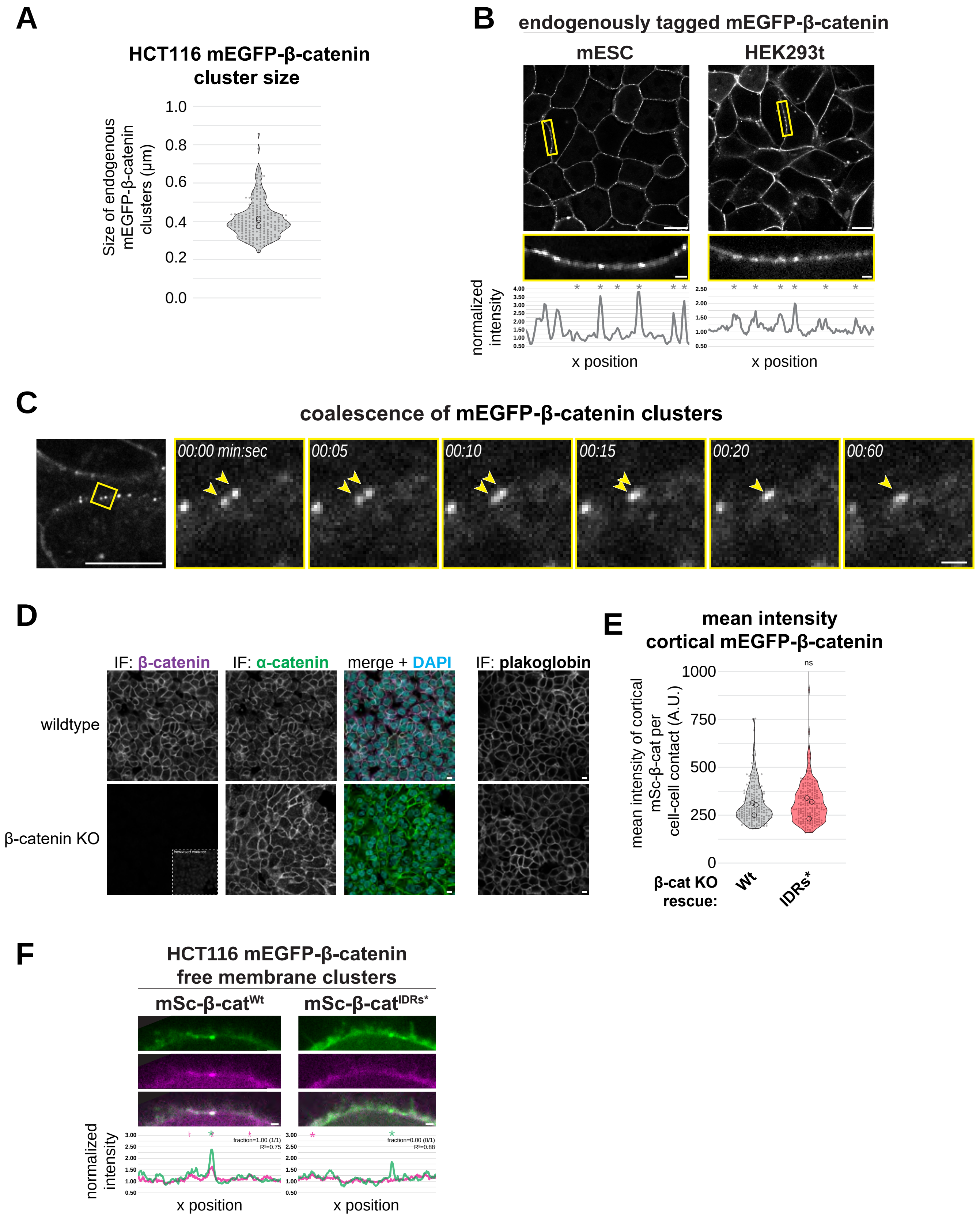


**Figure S2. Characterization of HCT116 cells with endogenously tagged, knock-out and addback of β-catenin.**
**A.** Quantification of the size of cortical mEGFP-β-catenin clusters in endogenously tagged HCT116 cells. Sizes are measured from line profiles of individual clusters as the full-width at half-maximum (see methods). Each point represents a measurement for a single cell-cell contact (n = 200), pooled from three independent measurements (medians indicated with circles). **B.** Representative confocal images of endogenously tagged mEGFP-β-catenin in mouse embryonic stem cells (mESC) and HEK293T cells, with zoom-ins (yellow box) of individual cell-cell contacts, showing that β-catenin organizes into small clusters at the cortex. Individual clusters are detected by peak detection on the corresponding line profile of the normalized fluorescent intensity of the junction, indicated by asterisks. See **Movies S5-6** for z-stacks of the displayed images. **C.** Representative still images (see **Movie S7**) of two cortical mEGFP-β-catenin clusters in HCT116 that coalesce and fuse into one cluster. An overview image (left) and the zoom-in of a cell-cell contact at different time frames (in min:sec) is shown, the coalescing clusters are highlighted with yellow arrowheads. **D.** Immunostainings of parental and β-catenin knock-out (KO) HCT116 cells for β-catenin (magenta) and α-catenin (green) together with DAPI (DNA; cyan), or plakoglobin. The absence of β-catenin does not result in the loss of cell-cell contacts (as indicated by the formation of a monolayer and the presence of junctional α-catenin), presumably due to compensation by the β-catenin homolog plakoglobin[74]. **E.** Quantification of the mean fluorescent intensity of β-catenin at cell-cell contacts in β-catenin KO HCT116 cells with addback of mSc-β-catenin$^{Wt}$ or mSc-β-catenin$^{IDRs*}$. Each point represents a measurement for a single cell-cell contact and correspond with the contacts measured in **Fig. 2E** (n = 181 and 186), pooled from three independent measurements (medians of each indicated with circles). ns = not significant; nested t-test. **F.** Representative confocal images of the free membrane of a HCT116 cell expressing both endogenously tagged mEGFP-β-catenin (green) and exogenously expressed mSc-β-cateninWt or mSc-β-cateninIDRs* (magenta). The corresponding line profiles show the normalized fluorescent intensity and detected peaks (asterisks) of both fluorophores. The fraction of peaks detected in the mEGFP channel with an overlapping peak in the mScarlet channel and the Pearson correlation (R2) between the line profiles of the two fluorophores are shown.Scale bars represent 10 µm (overview images) or 1 µm (zoom-ins of individual cell-cell contacts)

### Supplementary Figure 3

A

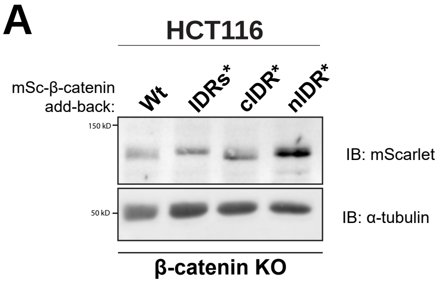


**Figure S3. Protein expression levels of β-catenin addback lines.**
**A.** Western blot of lysates of β-catenin knock-out (KO) HCT116 cells with addback of mSc-β-catenin$^{Wt}$, mSc-β-catenin$^{IDRs*}$, mSc-β-catenin$^{nIDR*}$ or mSc-β-catenin$^{cIDR*}$ and probed for mScarlet and α-tubulin. Note that the mScarlet antibody was used to compare the expression levels of ectopic wildtype and mutated β-catenin, excluding potential effects of the introduced mutations in β-catenin on antibody affinity.

### Supplementary Figure 4

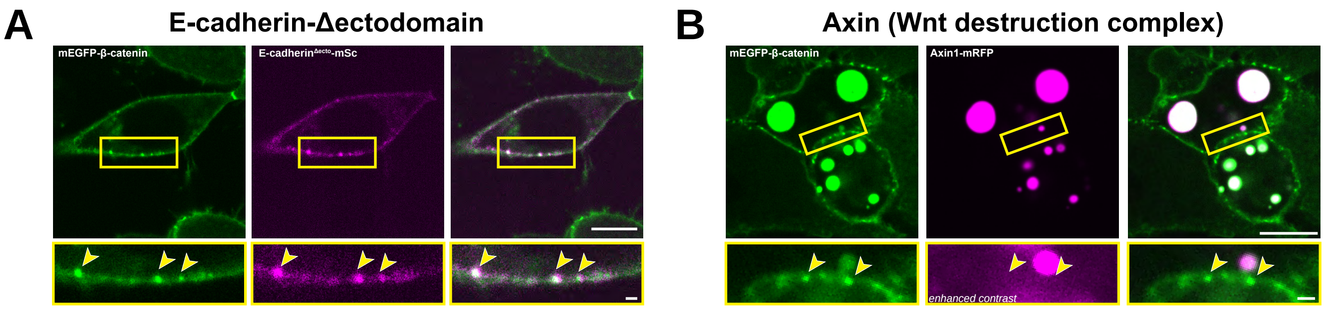


**Figure S4. Characterization of E-cadherin integration into IDR-dependent β-catenin clusters**
**A.**Representative confocal image and zoom-in (yellow box) of individual cell-cell contact of HCT116 cells expressing both endogenously tagged mEGFP-β-catenin (green) and E-cadherin$^{\Delta ecto}$ (magenta). This truncated version of E-cadherin still integrates in cortical β-catenin clusters at the free membrane, indicating E-cadherin cis-interactions between its extracellular domain are dispensable for cortical cluster formation in HCT116 cells. Note that cells overexpressing E-cadherin$^{\Delta ecto}$ were barely capable of forming cell-cell contacts due to their inability to form trans-interactions. **B.** Representative confocal image and zoom-in (yellow box) of individual cell-cell contact of HCT116 cells expressing both endogenously tagged mEGFP-β-catenin (green) and exogenously expressed Axin1-mRFP (magenta). Axin1 does not colocalize with mEGFP-β-catenin clusters at cell-cell contacts (indicated by yellow arrowheads), indicating that cortical β-catenin clusters do not represent clusters containing components of the destruction complex. Exogenous expression of Axin has been previously shown to form cytosolic biomolecular condensates[75], in which we find endogenous mEGFP-β-catenin to co-integrate.

### Supplementary Figure 5

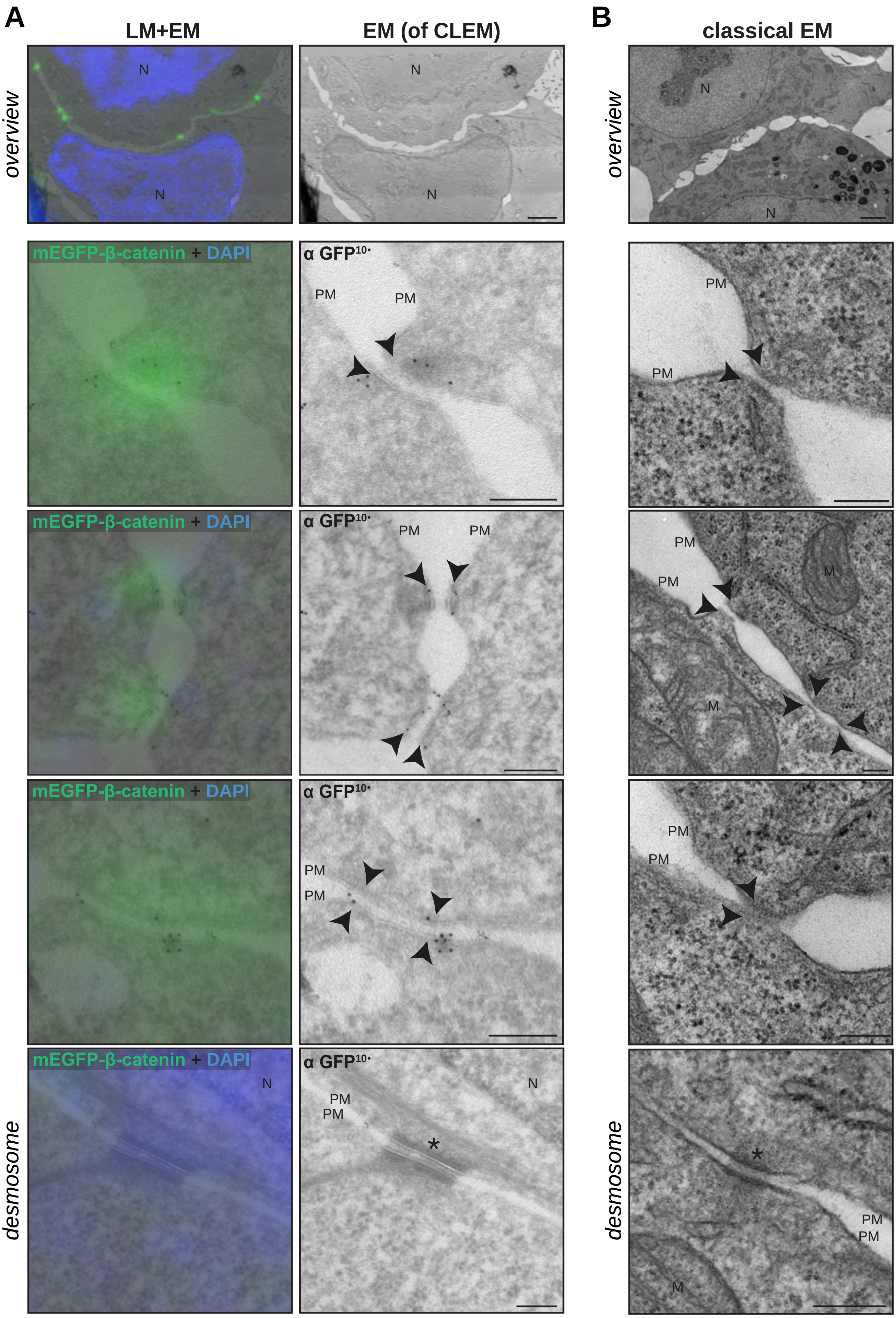

### Supplementary Figure 5 (continued)

C

disruption of the actin network

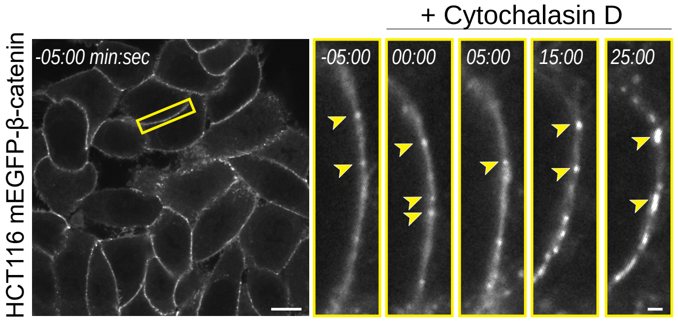


D

FRAP cell-cell cluster

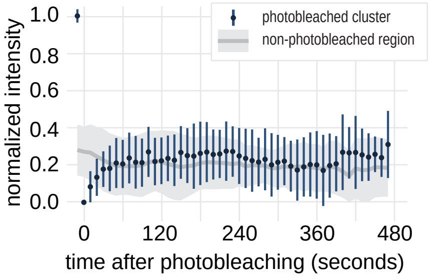


**Figure S5. Protein expression levels and monolayer formation in β-catenin knock-out and addback lines**

**A.**Additional Correlative Light and Electron Microscopy (CLEM) images of HCT116 cells with endogenously tagged mEGFP-β-catenin in 90-100 nm thawed cryosections (similar to **Fig. 5A**). Light-microscopy (LM) images of mEGFP-β-catenin (green) together with DAPI (blue) were overlayed with EM images (LM+EM). The sample also contains labeling of mEGFP with 10 nm gold particles (αGFP10). Three examples of cortical mEGFP-β-catenin clusters and an mEGFP-negative desmosome (marked *) are shown. Note cell-cell proximity and electron-dense clusters at sites of mEGFP-β-catenin localization (arrowheads). **B.** EM images of cell adhesion sites from resin-embedded HCT116 mEGFP-β-catenin cells. Note the focal nature of the cell-cell contacts and general lack of long, linear cell adhesions in these cells. We recognize two morphologically distinct types of cell-cell contact: one are likely desmosomes (bottom right image, marked *) as identified by prominent, electron-dense bands below the PM. The other appears as clear intercellular attachments but shows less pronounced intracellular protein density (arrowheads), resembling the β-catenin-positive sites observed in CLEM. **C.** Representative time-lapse images (see **Movie S8**) of endogenous mEGFP-β-catenin at a cell-cell contact (zoom-in; yellow box) in HCT116 cells upon the addition of 2 µg/ml Cytochalasin D. The cortical β-catenin clusters (marked by yellow arrowhead) increase in intensity and coalesce into bigger clusters over time. Time point 0 marks the addition of Cytochalasin D. **D.** Quantification of the normalized fluorescent signal (mean ± SD) before and after photobleaching of mEGFP-β-catenin clusters at cell-cell contacts (n = 15). Normalized fluorescent intensities of single clusters were measured every 10 seconds (blue dots). Intensities of a distant region of the junction was similarly measured in each frame (gray line ±SD), to show that the recovery of the clusters is similar to the level of the linear junction. Scale bars represent 2 µm (overview EM images A-B), 200 nm (EM zoom-ins; A-B), 10 µm (overview image C) or 1 µm (cell-cell contact; C). M, mitochondrion; N, nucleus; PM, plasma membrane; V, vesicle.

### Supplementary Figure 6

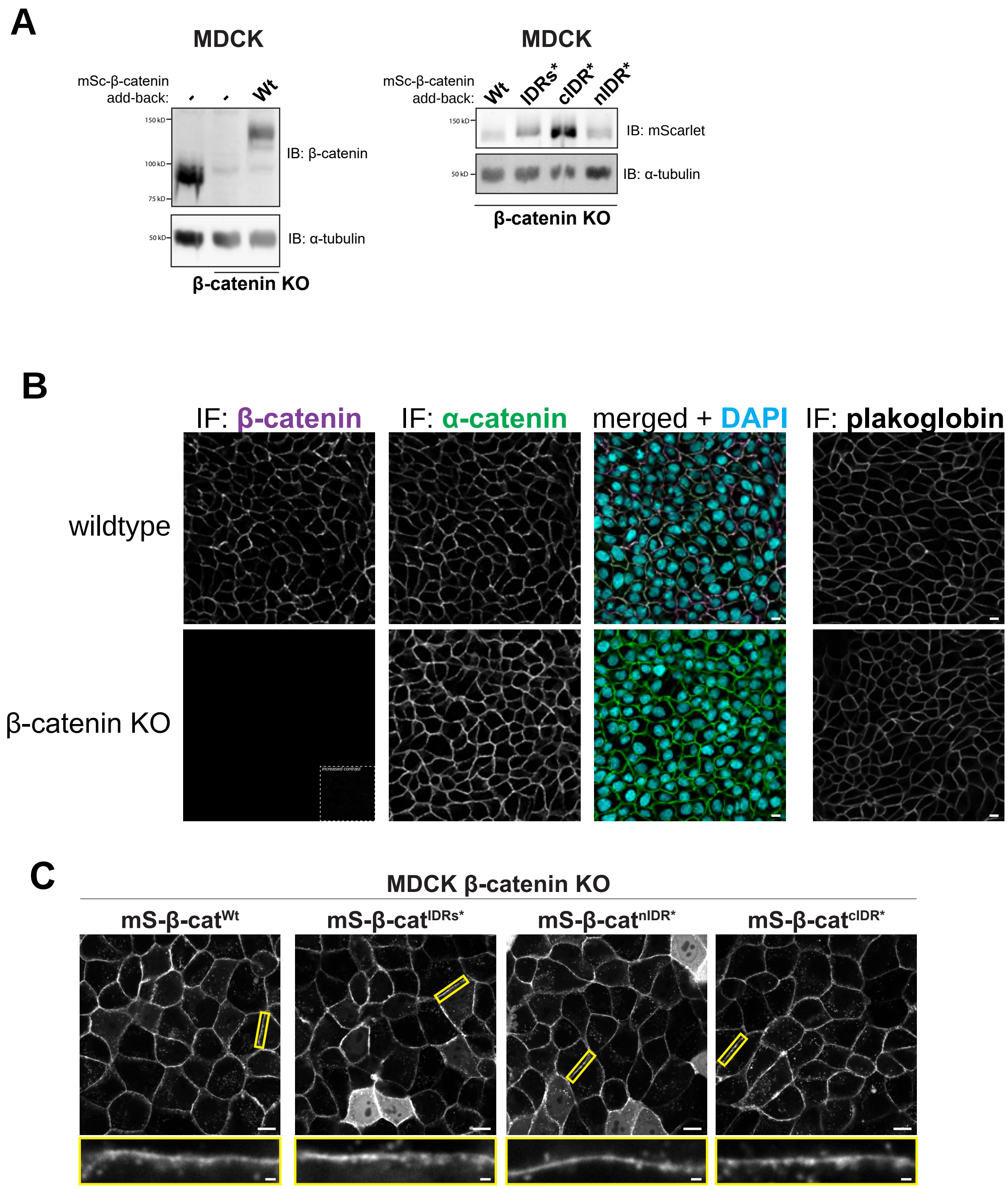


**Figure S6. Protein expression levels and monolayer formation in β-catenin knock-out and addback lines**

**A.** Left: Western blot of lysates of parental, β-catenin knock-out (KO) and addback with mSc-β-catenin$^{Wt}$ MDCK cells probed for β-catenin and α-tubulin. Right: Western blot of lysates of β-catenin KO MDCK cells with addback mSc-β-catenin$^{Wt}$, mSc-β-catenin$^{IDRs*}$, mSc-β-catenin$^{nIDR*}$ or mSc-β-catenin$^{cIDR*}$ probed for mScarlet and α-tubulin. Note that the mScarlet antibody was used to compare the expression levels of ectopic wildtype and mutated β-catenin, excluding potential effects of the introduced mutations in β-catenin on antibody affinity. **B.** Immunostainings of parental and β-catenin KO MDCK cells for β-catenin (magenta) and α-catenin (green) together with DAPI (DNA; cyan), or plakoglobin. The absence of β-catenin does not result in the loss of cell-cell contacts (as indicated by the formation of a monolayer and the presence of junctional α-catenin), presumably due to compensation by β-catenin homolog plakoglobin[73]. **C.** Representative examples of monolayers of β-catenin KO MDCK cells with addback of mSc-β-catenin$^{Wt}$, mSc-β-catenin$^{IDRs*}$, mSc-β-catenin$^{nIDR*}$ or mSc-β-catenin$^{cIDR*}$ probed for mScarlet and α-tubulin, 24 hours after seeding. Zoom-in (yellow box) shows individual cell-cell contact. While the efficiency of de novo contact formation is perturbed in IDR-mutant β-catenin lines (**Fig. 6**), when given sufficient time all mutant cells are able to establish cell-cell adhesion and develop into confluent epithelial monolayers. Furthermore, these lines have no noticeable differences in junction morphology compared to wildtype cells, in which the clusters observed during the initial formation of a novel cell-cell contact have reorganized into a uniform junction. Scale bars represent 10 µm (overview images) or 1 µm (zoom-ins of cell-cell contacts).

#### List of Supplementary Movies

**Movies S1 and S2** - Representative movies fusion events of droplets containing mEGFP-β-cat$^{Wt}$ (green), mCherry-α-cat (magenta) and E-cad$^{cyto}$-mTagBFP2 (cyan). The indicated proteins were premixed at a concentration of 1 μM before the addition of 10% PEG-8000. Time in minutes:seconds, scale bar represents 1 μm.

**Movie S3** - Representative confocal z-stack (0.2 μm interval, looped) of endogenously tagged mEGFP-b-catenin in HCT116 cells at a confluent density. Scale bar represents 10 μm.

**Movie S4** - Representative confocal z-stack (0.2 μm interval, looped) of endogenously tagged mEGFP-b-catenin in HCT116 cells at low density, showing clusters at the free membrane. Scale bar represents 10 μm.

**Movie S5** - Representative confocal z-stack (0.2 μm interval, looped) of endogenously tagged mEGFP-b-catenin in mouse embryonic stem cells (mESCs). Scale bar represents 10 μm.

**Movie S6** - Representative confocal z-stack (0.2 μm interval, looped) of endogenously tagged mEGFP-b-catenin in HEK293t cells. Scale bar represents 10 μm.

**Movie S7** - Representative time-lapse imaging of endogenously tagged mEGFP-b-catenin in HCT116 cells showing the coalescence of two clusters. Time in hours:minutes:seconds; scale bar represents 1 μm.

**Movie S8** - Representative time-lapse imaging of endogenously tagged mEGFP-b-catenin in HCT116 cells upon the addition of 2 μg/ml Cytochalasin D. Time in hours:minutes:seconds since the addition of Cytochalasin D; scale bar represents 10 μm.

**Movie S9** - Representative time-lapse imaging of Fluorescent Recovery After Photobleaching (FRAP) experiment. A single mEGFP-β-catenin cluster at a cell-cell contact of endogenously tagged HCT116 cells was photobleached (area indicated with yellow box, time point 0) and imaged over time. Time in hours:minutes:seconds; scale bar represents 1 μm.

**Movie S10** - Representative time-lapse imaging of Fluorescent Recovery After Photobleaching (FRAP) experiment. A single mEGFP-β-catenin cluster at the free membrane of endogenously tagged HCT116 cells was photobleached (area indicated with yellow box, time point 0) and imaged over time. Time in hours:minutes:seconds; scale bar represents 1 μm.

**Movie S11** - Representative time-lapse movie of normal *de novo* contact formation in β-catenin knock-out (KO) MDCK cells with mSc-β-catenin$^{Wt}$ addback (Fire LUT). Time in hours:minutes:seconds since the initial contact formation, scalebar represents 10 μm.

**Movie S12** - Representative time-lapse movie showing clustering defects in β-catenin knock-out (KO) MDCK cells with mSc-β-catenin$^{cIDR*}$ addback (Fire LUT) during *de novo* contact formation. Time in hours:minutes:seconds since the initial contact formation, scalebar represents 10 μm.

**Movie S13** - Representative time-lapse movie demonstrating a less efficient *de novo* contact formation in β-catenin knock-out (KO) MDCK cells with mSc-β-catenin$^{IDRs*}$ addback (Fire LUT). Both novel contacts show a reduced level of β-catenin clustering and a delay in cortical enrichment, whereas the right contact fails to enrich β-catenin entirely. Time in hours:minutes:seconds since the initial contact formation, scalebar represents 10 μm.

**Movie S14** - Representative time-lapse movie showing clustering defects and a failure to form a linear junction in β-catenin knock-out (KO) MDCK cells with mSc-β-catenin$^{nIDR*}$ addback (Fire LUT) during *de novo* contact formation. Time in hours:minutes:seconds since the initial contact formation, scalebar represents 10 μm.

**Movie S15** - Representative time-lapse movie of normal *de novo* contact formation in β-catenin knock-out (KO) MDCK cells with mSc-β-catenin$^{Wt}$ addback (Fire LUT). Time in hours:minutes:seconds since the initial contact formation, scalebar represents 10 μm.

**Movie S16** - Representative time-lapse movie showing a delay in enrichment in β-catenin knock-out (KO) MDCK cells with mSc-β-catenin$^{cIDR*}$ addback (Fire LUT) during *de novo* contact formation. Time in hours:minutes:seconds since the initial contact formation, scalebar represents 10 μm.

**Movie S17** - Representative time-lapse movie showing a failure to establish a linear cell-cell contact and instead maintain their dynamic protrusions in β-catenin knock-out (KO) MDCK cells with mSc-β-catenin$^{cIDR*}$ addback (Fire LUT). Time in hours:minutes:seconds since the initial contact formation, scalebar represents 10 μm.

**Movie S18** - Representative time-lapse movie showing an unsuccessful *de novo* contact formation and instead junction breakage in β-catenin knock-out (KO) MDCK cells with mSc-β-catenin$^{cIDR*}$ addback (Fire LUT). Time in hours:minutes:seconds since the initial contact formation, scalebar represents 10 μm.
